# Oxytocin promotes epicardial cell activation and heart regeneration after cardiac injury

**DOI:** 10.1101/2021.11.01.466355

**Authors:** Aaron H. Wasserman, Amanda R. Huang, Yonatan R. Lewis-Israeli, McKenna D. Dooley, Allison L. Mitchell, Manigandan Venkatesan, Aitor Aguirre

## Abstract

Cardiovascular disease (CVD) is one of the leading causes of mortality worldwide, and frequently leads to massive heart injury and the loss of billions of cardiac muscle cells and associated vasculature. Critical work in the last two decades demonstrated that these lost cells can be partially regenerated by the epicardium, the outermost mesothelial layer of the heart, in a process that highly recapitulates its role in heart development. Upon cardiac injury, mature epicardial cells activate and undergo an epithelial-mesenchymal transition (EMT) to form epicardial-derived progenitor cells (EpiPCs), multipotent progenitors that can differentiate into several important cardiac lineages, including cardiomyocytes and vascular cells. In mammals, this process alone is insufficient for significant regeneration, but it might be possible to prime it by administering specific reprogramming factors, leading to enhanced EpiPC function. Here, we show that oxytocin (OXT), a hypothalamic neuroendocrine peptide, induces epicardial cell proliferation, EMT, and migration in a model of human induced pluripotent stem cell (hiPSC)-derived epicardial cells. In addition, we demonstrate that OXT is produced after cardiac cryoinjury in zebrafish, and that it elicits significant epicardial activation promoting heart regeneration. Oxytocin signaling is also critical for proper epicardium development in zebrafish embryos. The above processes are significantly impaired when OXT signaling is inhibited chemically or genetically through RNA interference. RNA sequencing data suggests that the transforming growth factor beta (TGF-β) pathway is the primary mediator of OXT-induced epicardial activation. Our research reveals for the first time an evolutionary conserved brain-controlled mechanism inducing cellular reprogramming and regeneration of the injured mammalian and zebrafish heart, a finding that could contribute to translational advances for the treatment of cardiac injuries.

## INTRODUCTION

Cardiovascular disease (CVD) is the leading cause of mortality in the United States and the rest of the developed world (Virani et al., 2020). These diseases often lead to severe cardiac events, such as a myocardial infarction (MI), which can result in the death of more than 25% of cardiac myocytes, the chief contractile cells of the adult heart. If left untreated, this condition can lead to heart failure (Laflamme & Murry, 2011). Cardiomyocytes (CMs) are terminally differentiated cells, and as such, their innate regenerative capacity is extremely limited and insufficient to restore function to lost myocardium. Instead, the injured heart typically resorts to wound repair, replacing dead cells with fibrotic scar tissue (Bergmann et al., 2009; Fan et al., 2012). Therefore, the ability to induce heart regeneration, which results in growth and proliferation of endogenous diseased cells (and subsequent restoration of cardiac function), would be paramount in the discovery of new therapeutic strategies for the successful treatment of MI and other types of CVD (Laflamme & Murry, 2011; Uygur & Lee, 2016).

The epicardium is the outermost mesothelial layer of the heart that serves a structural and protective role in healthy individuals (Riley, 2012). While its activity level decreases as development proceeds, in adults, it becomes activated upon injury and recapitulates its embryonic phenotype (Peralta et al., 2014; Simões & Riley, 2018). Upon activation, a subset of epicardial cells migrate into the subepicardial space and undergo epithelial-mesenchymal transition (EMT) to form epicardial-derived progenitor cells (EpiPCs). These EpiPCs are multipotent cardiac progenitors (the only widely accepted source) that contribute to multiple different cell lineages in the heart and can aid in the healing and repair process (Quijada et al., 2020; Rao & Spees, 2017; Smits et al., 2018). While there is evidence to suggest that EpiPCs can differentiate into cardiomyocytes during development (Cai et al., 2008; Zhou et al., 2008), whether this process occurs in adult EpiPCs is a matter of ongoing debate. Adult epicardial cells also secrete paracrine factors to modulate heart repair and regeneration post-MI (Wei et al., 2015; Zhou et al., 2011). Therefore, the activated epicardium might be the key to unlocking cardiac regeneration pathways in humans.

The brain plays a central role in several energetically demanding processes that serve to maintain body homeostasis (Dampney, 2016; Guyenet & Bayliss, 2015; Morrison, 2016; Roh et al., 2016). In addition, the brain-heart signaling axis regulates heart rate, blood pressure, and systolic and diastolic function (Dampney, 2016; Palma & Benarroch, 2014). The neural endocrine structures, such as the hypothalamus and pituitary, play a particularly important role in maintaining normal cardiovascular function through the release of various hormones (Gordan et al., 2015; Rhee & Pearce, 2011). Therefore, it stands to reason that a process as biologically and energetically demanding as heart regeneration does not happen cell-autonomously and might be under central hormonal control. In fact, there is evidence that several hormones such as estrogen (Xu et al., 2020), thyroid hormones (Hirose et al., 2019), and cortisol (Sallin & Jaźwińska, 2016) affect cardiac regeneration. In addition, a recent study indicates that damage to the hypothalamus inhibits endogenous regenerative processes in vertebrates (Zhang et al., 2018). These findings, combined with the fact that injury-induced epicardial activation can be primed by specific signaling factors (Smart et al., 2011), led us to hypothesize that after cardiac injury, one or more neuroendocrine hormones are released from the hypothalamus into the bloodstream to facilitate heart regeneration. Proper identification and characterization of these hormones will be critical in advancing the translational potential of cardiac regeneration research.

Here, we present evidence that oxytocin (OXT), a nine amino acid neuropeptide produced by the hypothalamus and released by the posterior pituitary, may be the missing link in achieving sufficient EpiPC proliferation, activation, and migration to fully regenerate the lost myocardium in the injured heart. We utilize both *in vitro* and *in vivo* injury models to identify and describe the direct involvement of OXT in these processes. First, we differentiate and characterize a mature-like model of human iPSC-derived epicardial cells (hEpiCs) and show that the peptide induces proliferation and activation in these cells by acting through its receptor (OXTR). We then show that oxytocin is released from the brain after cardiac cryoinjury in zebrafish, a process that is critical for heart regeneration and epicardial activation. In addition, inhibition of OXT signaling delays regeneration of adult hearts and epicardium formation in zebrafish embryos. RNA sequencing (RNA-seq) analyses suggest that the TGF-β pathway is the primary mediator of OXT-induced epicardial activation. The findings of this study reveal a novel role for the brain-heart signaling axis in regulating cardiac regeneration and elucidate the critical importance of the epicardium in this process.

## MATERIALS AND METHODS

### Stem cell culture

Human iPSCs (hiPSC-L1) were cultured on 6 well plates coated with growth factor-reduced Matrigel (Corning) in an incubator at 37°C, 5% CO_2_. Stem cells were maintained in Essential 8 Flex medium (E8 Flex; Thermo Fisher Scientific) containing 1% penicillin/streptomycin until ~70% confluency was reached, at which point cells were split into new wells using ReLeSR passaging reagent (Stem Cell Technologies) with 2 μM ROCK inhibitor (Thiazovivin/TZV; Selleck Chemicals) added to prevent cells from undergoing apoptosis while in suspension. All hiPSC lines were periodically validated for pluripotency and genomic stability.

### hiPSC differentiation to hEpiCs

hiPSCs were differentiated to epicardial monolayers using a modified version of a previously described protocol (Bao et al., 2016, 2017). Cells were dissociated using Accutase (Innovative Cell Technologies), re-plated on Matrigel coated plates, and incubated overnight in E8 Flex + 2 μM TZV. Once hiPSC monolayers reached ~90% confluency, epicardial differentiation was started via the addition of 12 μM CHIR99021 (Selleck Chemicals) in RPMI/B-27 minus insulin (Gibco; Day 0) for 24 hours. On Day 3, cells were exposed to 2 μM Wnt-C59 (Selleck Chemicals) in RPMI/B-27 minus insulin for 48 hours. On Day 6, cardiac progenitors were then re-plated in RPMI/B-27 (Gibco) with 100 μg/mL Vitamin C (Vit. C) using Accutase and TZV. Cells were directed down the epicardial lineage by the consecutive addition of 9 μM CHIR99021 on Days 7 and 8 and maintained in RPMI/B-27/Vit. C until passage. On day 12, hEpiCs were again re-plated using Accutase and TZV and 2 μM SB431542 (TGF-β receptor inhibitor; Selleck Chemicals) was added to the media to prevent cells from spontaneously undergoing EMT. Epicardial cells were cultured long-term in RPMI/B-27/Vit. C + 2 μM SB431542 and split when full confluency was reached. To ensure that hEpiCs were as mature as possible for experiments, Vitamin C and SB431542 were removed from the culture system 5 days prior to experiments. OXT and other compounds were added directly to the media 2 days later, for a total exposure time of 3 days, unless otherwise indicated. Automated cell counting was conducted using the Cytation 3 and Cytation 5 Cell Imaging Multi-Mode Readers (Biotek).

### hiPSC lentiviral transduction

Bacteria carrying the plasmid for shRNA-mediated knockdown of *OXTR* and a scrambled plasmid (both designed with VectorBuilder) were grown on LB agar plates and isolated colonies were expanded in LB broth, both containing ampicillin. The plasmids carried an ampicillin resistance cassette, allowing for their growth. Plasmid DNA was isolated from the bacteria by midiprep (Zymo Research), and purified DNA was transfected into 40% confluent HEK293T cells using Lipofectamine (Invitrogen). Lentiviral packaging plasmids (pVSVg, psPAX2) were also transfected at this time, thereby allowing the generation of a functional lentivirus containing the shRNA molecules of interest. Viral supernatant was collected and concentrated after 48 hours and transduced directly into hiPSCs at low to mid-confluency along with 8 μg/mL polybrene (Fisher Scientific). Because all plasmids contained puromycin-resistance cassettes, the stem cell media was changed the next day to E8 Flex containing 0.5 μg/mL puromycin (Thermo Fisher Scientific) to facilitate colony selection. hiPSCs were maintained in puromycin for 5 days and surviving monoclonal colonies were re-plated and expanded to generate new hiPSC lines.

### Zebrafish cardiac cryoinjury

Adult zebrafish hearts were subjected to cardiac cryoinjury as described previously (González-Rosa et al., 2011; González-Rosa & Mercader, 2012). Briefly, fish were anesthetized with 0.65 mM tricaine (MS-222) and placed on a wet sponge ventral side up under a stereomicroscope (Leica). The chest cavity was opened by cutting through the pericardium until the point where the beating heart was clearly visible. For cryoinjury, a liquid nitrogen cooled metal probe was gently placed on the apex of the ventricle for ~45 seconds. Sham operations were carried out by simply opening the chest cavity without injury. Fish were closely monitored for the first 2 hours after surgery and twice daily thereafter until ready for organ collection. Pharmacological inhibition of oxytocin signaling was achieved by injecting 2-3 μL of water or 1 μM L-368,899 (a non-peptide OXTR antagonist) directly into the intrathoracic cavity, as described previously (Bise & Jaźwińska, 2019). Initial injections were administered 24 hours after cryoinjury with repeat injections every other day throughout the experiment. All zebrafish were maintained in a dedicated facility between 27-28°C on a 14-hour light/10-hour dark cycle. Equal numbers of male and female fish were used for cryoinjuries.

### Zebrafish embryo experiments

One female and one male adult fish were placed in specialized breeding tanks with a partition between them the night before experiments. The partition was removed early the next morning and fish were given ~15 minutes to breed. Fertilized embryos were collected and placed in embryo medium containing 4.96 mM NaCl, 0.179 mM KCl, 0.329 mM CaCl_2_ • 2H_2_O, and 0.401 mM MgCl_2_ • 6H_2_O dissolved in water. The appropriate concentration of atosiban (competitive OXTR antagonist) was added directly to the medium and embryos were stored in an incubator at 28°C for the duration of the experiment. Dead embryos and discarded chorions were removed daily. Developing fish were imaged at 1, 2, 3, 5, and 7 days post-fertilization (dpf) with the Leica M165 FC stereomicroscope with a DFC7000 T fluorescence camera so that epicardium and myocardium formation could be observed in real time. Epicardial cell number was counted manually at 3, 5, and 7 dpf.

### Gene expression analysis

Total RNA was extracted from samples using the RNeasy Mini Kit (Qiagen). Cells were lysed directly in their culture wells and tissues were lysed and homogenized using the Bead Mill 4 Homogenizer (Fisher Scientific). Once extracted, RNA was quantified using a NanoDrop (Mettler Toledo), with a concentration of at least 10 ng/μL being required to proceed with reverse transcription. cDNA was synthesized using the Quantitect Reverse Transcription Kit (Qiagen) and stored at −20°C for further use. Primers for qRT-PCR were designed using the Primer Quest tool (Integrated DNA Technologies) and SYBR Green (Qiagen) was used as the DNA intercalating dye. qRT-PCR plates were run using the QuantStudio 5 Real-Time PCR system (Applied Biosystems) with a total reaction volume of 20 μL. Expression levels of genes of interest were normalized to *HPRT1* levels and fold change values were obtained using the 2^-ΔΔCT^ method. At least 3-5 independent samples were run for each gene expression assay.

### Immunofluorescence

hEpiC samples were first transferred onto Millicell EZ slides (MilliporeSigma), fixed in 4% paraformaldehyde (PFA) solution, and washed with PBS + 1.5 g/L glycine. Zebrafish tissue samples were extracted, washed in PBS + 0.5 mM EDTA solution, and fixed in 4% PFA for one hour. After fixation, organs were washed with PBS-glycine and transferred into a PBS + 30% sucrose solution for at least 48 hours. Samples were then embedded in Optimal Cutting Temperature (OCT) compound (Electron Microscopy Sciences), frozen in a mold, and sectioned at 10 μm thickness using the Leica CM3050 S cryostat. Sectioned tissue was stored at 4°C for further use. Once ready for staining, antigen retrieval was performed by immersing slides in a sodium citrate buffer (10 mM sodium citrate, 0.05% Tween 20, pH 6.0) for ~45 minutes at 95-100°C. All samples (cells and tissues) were blocked and permeabilized with 10% normal donkey serum, 0.5% bovine serum albumin (BSA), and 0.5% Triton X-100 in PBS for 1 hour at room temperature. After washing, primary antibodies were diluted in antibody solution (1% normal donkey serum, 0.5% BSA, 0.5% Triton X-100 for tissues, 0.05% Triton X-100 for cells, all dissolved in PBS), added to the samples, and incubated overnight at 4°C. The next day, cells and tissues were washed, and secondary antibodies were diluted in antibody solution (with 0.05% Triton X-100) and added for 2 hours at room temperature. 4’,6-diamidino-2-phenylindole (DAPI, Thermo Fisher Scientific) was then added immediately at a concentration of 1:1000 to label DNA. Stained slides were washed 3 times in PBS and No.1 coverslips (VWR) were added using ProLong Gold Antifade Mountant (Thermo Fisher Scientific).

### Histology

Masson’s trichrome staining was carried out using a kit from IMEB Inc. and following manufacturer’s instructions. Briefly, sectioned tissue was immersed in Bouin’s solution for ~50 minutes at 56-64°C to improve staining quality and washed in tap water until the yellow color was fully removed from the hearts. Slides were then stained with Biebrich’s scarlet/acid fuchsin solution for 7 minutes, washed in deionized (DI) H_2_O, and immersed in phosphomolybdic/phosphotungstic acid solution for 18 minutes to decolorize fibrous tissue. Slides were then transferred directly to aniline blue solution for 12 minutes to stain collagen fibers, washed in DI H_2_O, and immersed in 1% acetic acid for 3 minutes to stabilize the staining. After washing, tissue was dehydrated by successive exposure to 95% ethanol, 100% ethanol, and xylene and coverslips were added using Eukitt quick-hardening mounting medium (Sigma-Aldrich). Scar size was quantified manually by tracing the injured myocardial area in ImageJ and dividing it by the total myocardial area. At least 2 sections from at least 3 independent hearts were analyzed for each condition at each time point.

### Confocal microscopy and image analysis

All samples were imaged using the Zeiss LSM880 NLO Confocal Microscope system or the Nikon A1 Confocal Laser Microscope. Images were analyzed and prepared for publication using Fiji software. Cell counts were performed manually using the “Wand” tool and automatically using the “Threshold” and “Analyze Particles” tools. Individuals who conducted manual cell counting of hEpiCs were blinded to experimental conditions and at least 5 images were analyzed for each group. For analysis of zebrafish confocal images, at least 3 independent fish from each condition were analyzed. Epicardial cell number was quantified by counting the number of GFP+ cells on the surface of the heart and in the subepicardial space per field of view. To ensure consistency in analysis, each image was taken immediately adjacent to the injured area. The number of h3p+/wt1b+ cells in the epicardial region was normalized to the number of DAPI-stained nuclei. Cardiomyocyte proliferation was quantified by manually counting the number of PCNA+ cells per field of view in the myocardium of the infarct border zone. At least two regions per heart were analyzed. Vascularization was quantified by measuring the GFP+ area within the injury region and dividing that value by the total injured area. In all cases, TNNT2 was utilized as a counterstain to allow for visualization of the cryoinjury. Quantification analyses were performed in a blinded fashion.

### RNA sequencing

RNA was extracted from 3 control and 3 hEpiC samples treated with 100 nM OXT as described above. RNA was quantified using a Qubit Fluorometer (Thermo Fisher Scientific) and samples were sent to the MSU Genomics core, where their quality was tested using the Agilent 2100 Bioanalyzer. Samples were sequenced using an Illumina HiSeq 4000. For RNA-seq sample processing, a pipeline was created in Galaxy. Briefly, sample run quality was assessed with FASTQC and alignment to hg38 was carried out using HISAT2. Counts were obtained using featureCounts and differential expression analysis was performed with EdgeR. Further downstream bioinformatic analysis was performed using Phantasus 1.11.0 (artyomovlab.wustl.edu/phantasus), ToppGene Suite (http://toppgene.cchmc.org), and Enrichr (https://maayanlab.cloud/Enrichr/).

### Statistical analysis

All graphs were made using GraphPad software. Statistical significance was evaluated with a standard unpaired Student’s t-test (2-tailed) when comparing two groups or with a 1-way ANOVA with Tukey’s or Dunnett’s post-test for multiple comparison analysis. Regardless of the test used, a P-value of less than 0.05 was considered statistically significant. All data are presented as mean ± SD or SEM and represent a minimum of 3 independent experiments with at least 3 technical replicates unless otherwise stated. For RNA-seq analysis, a false discovery rate (FDR) was used to determine statistical significance.

### Data availability

All RNA-seq datasets are deposited in Gene Expression Omnibus under GSE199427.

## RESULTS

### Establishing human iPSC-derived epicardial progenitor and mature-like epicardial cell cultures

To produce mature-like epicardial cells from human iPSCs (hEpiCs) we started by deriving epicardial progenitor cells using a 3-step Wnt modulation strategy (Bao et al., 2016, 2017) (**Figure 1A**). After 12 days of culture, hEpiCs display classic epithelial morphology (**Figure 1B**), express epicardial progenitor markers WT1 and TCF21 (**Figure 1C, D**), and show increased *WT1* and *CDH1* (epithelial cadherin) levels as they age (**Figure 1E, F**), suggesting an increase in epicardial nature. These cells can be routinely maintained and cultured in medium containing Vitamin C and SB431542 (a TGF-β receptor inhibitor), allowing them to retain their epithelial morphology for at least 20 passages; however, we found that these compounds prevent hEpiC maturation. As the epicardium develops and matures, its resident cells transition from a highly proliferative, migratory, and EMT state to a more mature, quiescent phenotype. In the mouse heart, key genes involved in epicardial activation are barely detectable by 3 months of age (Smart et al., 2011; Smits et al., 2018; Zhou et al., 2011). We hypothesized that removing Vitamin C and SB431542 from the hEpiC media for a short time would allow the cells to mature further in the epicardial lineage providing a better model to study epicardial activation *in vitro*. Thus, we removed these two factors and cultured hEpiCs for 5 days at confluence without passaging. Cells were then assayed for gene expression of common epicardial, smooth muscle, and fibroblast markers (to assess potential transdifferentiation into other cell types). Nearly 100% of hEpiCs maintained *WT1* and *TJP1* expression (the highly specific epithelial tight junction protein ZO-1) (**Figure 1G**), and no significant difference in expression of smooth muscle and fibroblast markers *CNN1, VIM*, and *ACTA2* was observed, suggesting maturated hEpiCs did not undergo EMT (**Figure 1H**). We found that maturation conditions reduced hEpiC proliferation (cell counts were 2-fold higher when cultured in media containing Vitamin C and SB431542 compared to when they were absent) (**Figure 1I**) and decreased progenitor and EMT marker expression (*WT1*, *TCF21, NT5E*), while at the same time increasing mature epithelial marker expression, such as *CDH1* (**Figure 1J**), confirming our hypothesis. Therefore, all experiments with epicardial cells were performed using this final media formulation to ensure that hEpiCs resembled epicardial cells in the adult heart as much as possible.

**Figure 1:**
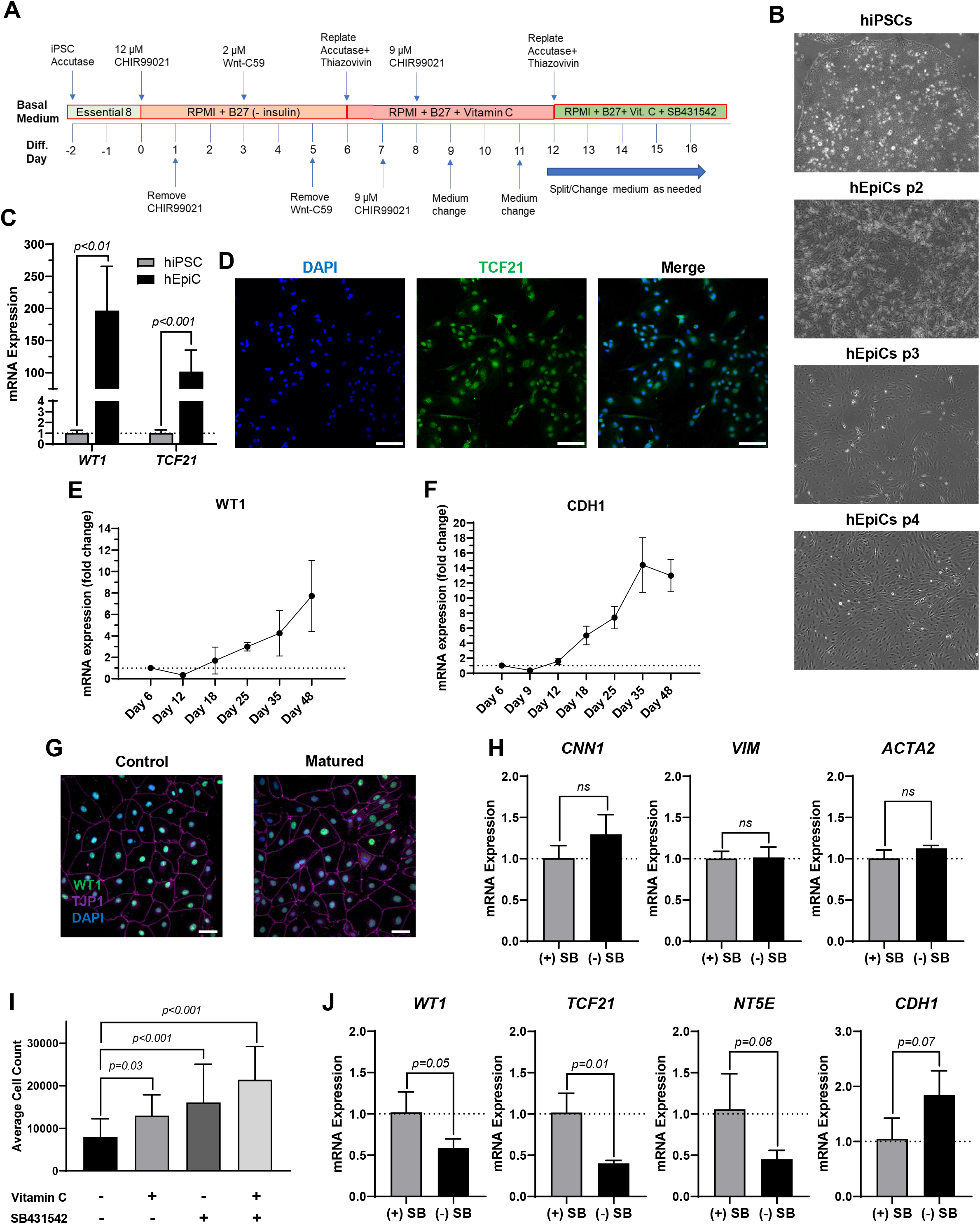
Differentiation of mature-like human epicardial cells from hiPSCs. **A)** Schematic of protocol used for hEpiC differentiation. **B)** Brightfield images of hEpiCs through four passages showing the gradual accumulation of the classic epithelial cell cobblestone morphology. **C)** qRT-PCR data for epicardial markers *WT1* and *TCF21* in hiPSCs and hEpiCs; n=4-6 per cell line. **D)** Confocal immunofluorescent images for TCF21 (green) and DAPI (blue) showing nearly 100% epicardial differentiation efficiency from hiPSCs; scale bar: 100 μm. **E-F)** Time course qRT-PCR data for *WT1* (E) and *CDH1* (F) throughout hEpiC differentiation, suggesting an increase in epicardial nature over time; n=3 per time point. **G)** Confocal immunofluorescent images showing robust expression of epicardial markers after removal of Vitamin C and SB431542 from culture media. Epicardial cells are labeled with WT1 (green), epithelial membranes with TJP1 (magenta), nuclei with DAPI (blue); scale bar: 50 μm. **H)** qRT-PCR data for hEpiCs in the presence or absence of SB431542, showing no change in smooth muscle or fibroblast differentiation; n=3 per condition. **I)** Absolute counts of DAPI-labeled nuclei after exposure to different combinations of Vitamin C and SB431542; n>20 per condition. **J)** qRT-PCR data for hEpiCs in the presence or absence of SB431542 (SB), showing an increase in maturity when this compound is removed from the media; n=3 per condition.

### Oxytocin induces human epicardial cell activation to a progenitor-like state

Because the brain is critical for maintenance of body homeostasis (Dampney, 2016; Guyenet & Bayliss, 2015; Morrison, 2016; Roh et al., 2016) and damage to the hypothalamus inhibits limb regeneration in vertebrates (Zhang et al., 2018), we hypothesized that the brain neuroendocrine system could also be involved in the process of heart regeneration. To identify potential neuroendocrine hormones involved in epicardial activation and possibly heart regeneration, we screened fifteen endogenous candidates in hEpiCs meeting the following two main criteria: 1) being involved in the physiological response to injury and 2) being produced and secreted by neuroendocrine tissue (see **Supplementary Table 1**). We identified 15 candidates and screened each of them for their proliferative effects by adding them to mature hEpiCs in culture. After 3 days, the percentage of phosphorylated histone H3 (H3P) positive cells was measured as a putative marker of proliferation using an high-content imaging microscope. Oxytocin (OXT), a neuroendocrine peptide released by the neurohypophysis, had the strongest proliferative effects (~3-fold versus control) and was selected for further downstream analysis (**Figure 2A**). We then exposed hEpiCs to different OXT concentrations in the physiological range and assessed cellular proliferation levels by two alternative methods, Ki67 staining and automated direct cell number counts using a nuclear dye (DAPI), as epicardial cells are mononucleated. We found that the percentage of hEpiCs double-positive for both WT1 and Ki67 increased by ~50% three days after addition of OXT (**Figure 2B, C**). Automated direct cell counting using DAPI staining produced similar results and indicated that the absolute number of epicardial cells doubled with oxytocin exposure (**Figure 2D**). We compared the effect of OXT to that of thymosin β4, a short peptide that has been shown to elicit strong epicardial activation *in vitro* and *in vivo* in mice before (Smart et al., 2007, 2011; Y. L. Wang et al., 2021). OXT caused a significantly more potent proliferative effect than thymosin β4 (**Figure 2E**). OXT also induced hEpiC epithelial-to-mesenchymal transition (EMT), as determined by an increase in gene expression for EMT markers *WT1*, *TCF21*, and *SNAI1* (**Figure 2F**). Taken together, these data suggest that oxytocin induces epicardial cell activation to a progenitor-like state.

**Figure 2:**
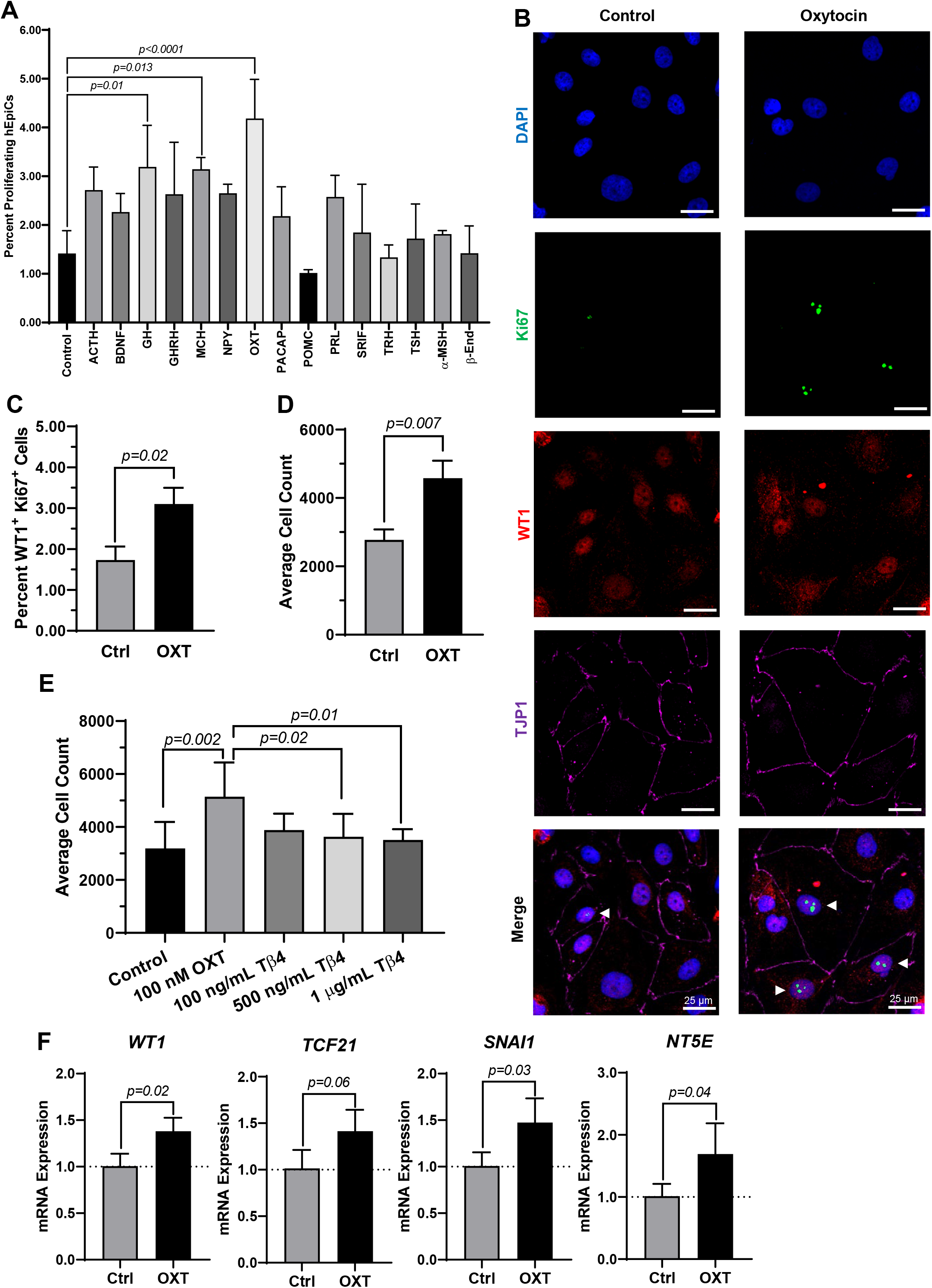
Oxytocin induces proliferation and activation of hiPSC-derived epicardial cells. **A)** Screening results for 15 candidate neuroendocrine peptides, shown as the percentage of hEpiCs expressing H3P after exposure to each compound n=3 per condition. **B-C)** Confocal immunofluorescent images (B) and quantification (C) of proliferating hEpiCs after 3 day OXT exposure. In (B), epicardial cells are labeled with WT1 (red), epithelial membranes are labeled with TJP1 (magenta), proliferating cells (arrowheads) are labeled with Ki67 (green), nuclei are labeled with DAPI (blue); n=8 samples per condition, scale bar: 25 μm. **D-E)** Absolute counts of DAPI-labeled epicardial cell nuclei after 3-day exposure to OXT (D) or thymosin β4 (E), a compound previously shown to induce epicardial activation; n≥8 per condition. **F)** qRT-PCR data for epicardial cells exposed to OXT, showing an increase in EpiPC (*WT1*, *TCF21*), EMT (*SNAI1*), and mesenchymal (*NT5E*) markers; n=4 per condition.

### Oxytocin exerts its pro-regenerative effects through the oxytocin receptor (OXTR)

We sought to gain some mechanistic insight as to how OXT causes its effects *in vitro*. There is only one oxytocin receptor in humans (OXTR), and it is a G-protein coupled receptor (GPCR) (Gimpl & Fahrenholz, 2001). We first conducted a dose-response experiment and quantified epicardial cell proliferation at each concentration. OXT had a concentration-dependent effect on hEpiC proliferation, with a maximal response (>2-fold increase in epicardial cell counts) occurring at 100 nM and an EC_50_ at ~300 pM (**Figure 3A and Supplementary Figure 1A**), consistent with a GPCR receptor-mediated response. Gene expression analysis showed that *OXTR* expression increased 20 to 30-fold in hEpiCs compared to undifferentiated hiPSCs (**Figure 3B**). Immunostaining and confocal imaging confirmed that OXTR is present on the cell membrane and in the cytoplasm of hEpiCs (**Figure 3C**). Epicardial cells did not express any detectable levels of *OXT* as assessed by qRT-PCR (data not shown). In addition to binding its own receptor, oxytocin can also bind to the arginine vasopressin (AVP) receptors in humans, albeit with lower affinity than to OXTR (Gimpl & Fahrenholz, 2001), however hEpiCs also did not express any of the three AVP receptor isoforms (data not shown). These data led us to believe that oxytocin acts through OXTR to induce epicardial activation. We created a knockdown *OXTR* hiPSC line using an shRNA-lentivirus targeting *OXTR*. This procedure resulted in a >90% knockdown in OXTR expression compared to scrambled epicardial cells (**Supplementary Figure 1B**) and did not affect the ability of hiPSCs to differentiate into hEpiCs (**Supplementary Figure 1C**). Exposure to 100 nM OXT led to strong hEpiC activation in scrambled cells, with an even greater response to LIT-001, a recently characterized and very potent non-peptide OXTR agonist (Frantz et al., 2018; Hilfiger et al., 2020) (**Supplementary Figure 1D**). These findings were confirmed when we stained the cells for proliferation marker Ki67. We observed a ~40% increase in proliferating hEpiCs in the scrambled line when challenged with OXT, but no response in the *shOXTR* line (**Figure 3D, E**). We next carried out qRT-PCR gene expression analyses to determine if EMT was also affected by *OXTR* knockdown. We observed a 2 to 4-fold increase in hEpiC activation (*WT1, TCF21*, and *SNAI1* marker expression) in scrambled cells, but these effects were entirely prevented in OXTR knocked-down hEpiCs (**Figure 3F, G**). We concluded that oxytocin acts through the OXTR to induce human epicardial cell activation *in vitro*.

**Figure 3:**
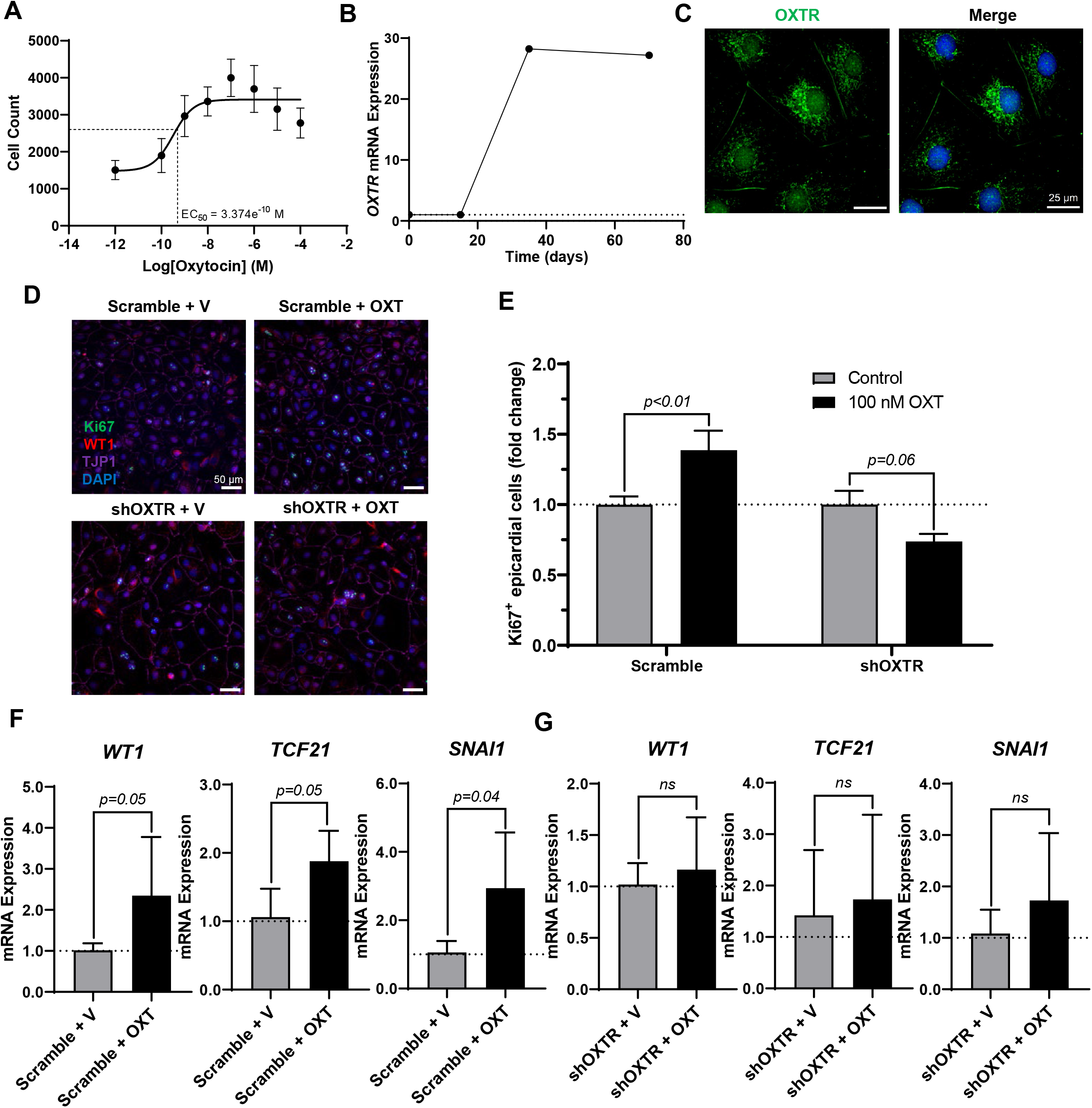
*OXTR* knockdown decreases proliferation and activation of hiPSC-derived epicardial cells. **A)** Dose-response data for hEpiCs exposed to different concentrations of OXT over the course of 3 days, expressed as number of nuclei at each concentration; n=10 per concentration. **B)** Time course qRT-PCR data for *OXTR* in hEpiCs. **C)** Confocal immunofluorescent images for OXTR (green) and DAPI (blue) showing oxytocin receptor expression on the cell membrane and in the peri-nuclear region; scale bar: 25 μm. **D-E)** Confocal immunofluorescent images (D) and quantification (E) of proliferating hEpiCs in both cell lines in the presence and absence of OXT at day 25 of differentiation. In (D), epicardial cells are labeled with WT1 (red), epithelial membranes are labeled with TJP1 (magenta), proliferating cells are labeled with Ki67 (green), nuclei are labeled with DAPI (blue); n=10 images per condition, scale bar: 50 μm. **F-G)** qRT-PCR data for scrambled (F) and shOXTR (G) hEpiCs, showing an increase in epicardial activation in control cells that is prevented after *OXTR* knockdown; n=6 per condition, V: Vehicle.

### Oxytocin acts through the TGF-β signaling pathway to induce epicardial activation

To obtain additional clues related to the potential mechanism of action driving the effects of oxytocin, we performed RNA-seq on hEpiCs in control and OXT-treated conditions as described above. RNA was collected for transcriptomic differential gene expression analysis after 3 days of OXT exposure. We found that oxytocin induced significant widespread gene expression changes (**Figure 4A**). Computational analysis using gene ontology identified upregulated and downregulated clusters consistent with our previous observations on the effects of oxytocin (induction of a progenitor-like state, increased proliferation, EMT) (**Figure 4B, C**). Of particular interest was the upregulation of TGF-β/BMP pathway biological processes, as well as developmental ones (**Figure 4B**). Among downregulated processes, a series of metabolic functions well-ascribed to the epicardium were identified, consistent with our proposed model of mature epicardial cells becoming epicardial progenitors upon OXT treatment. Leading genes driving the TGF-β pathway activation were ligands *LEFTY2, GDF15*, and *INHBB*, and the BMP pathway activator *TMEM100* (**Figure 4D**). Overall, these data suggest that oxytocin, through the activation of OXTR, can promote the expression of proteins involved in TGF-β and BMP signaling in the epicardium, leading to increased epicardial progenitor cell pools.

**Figure 4:**
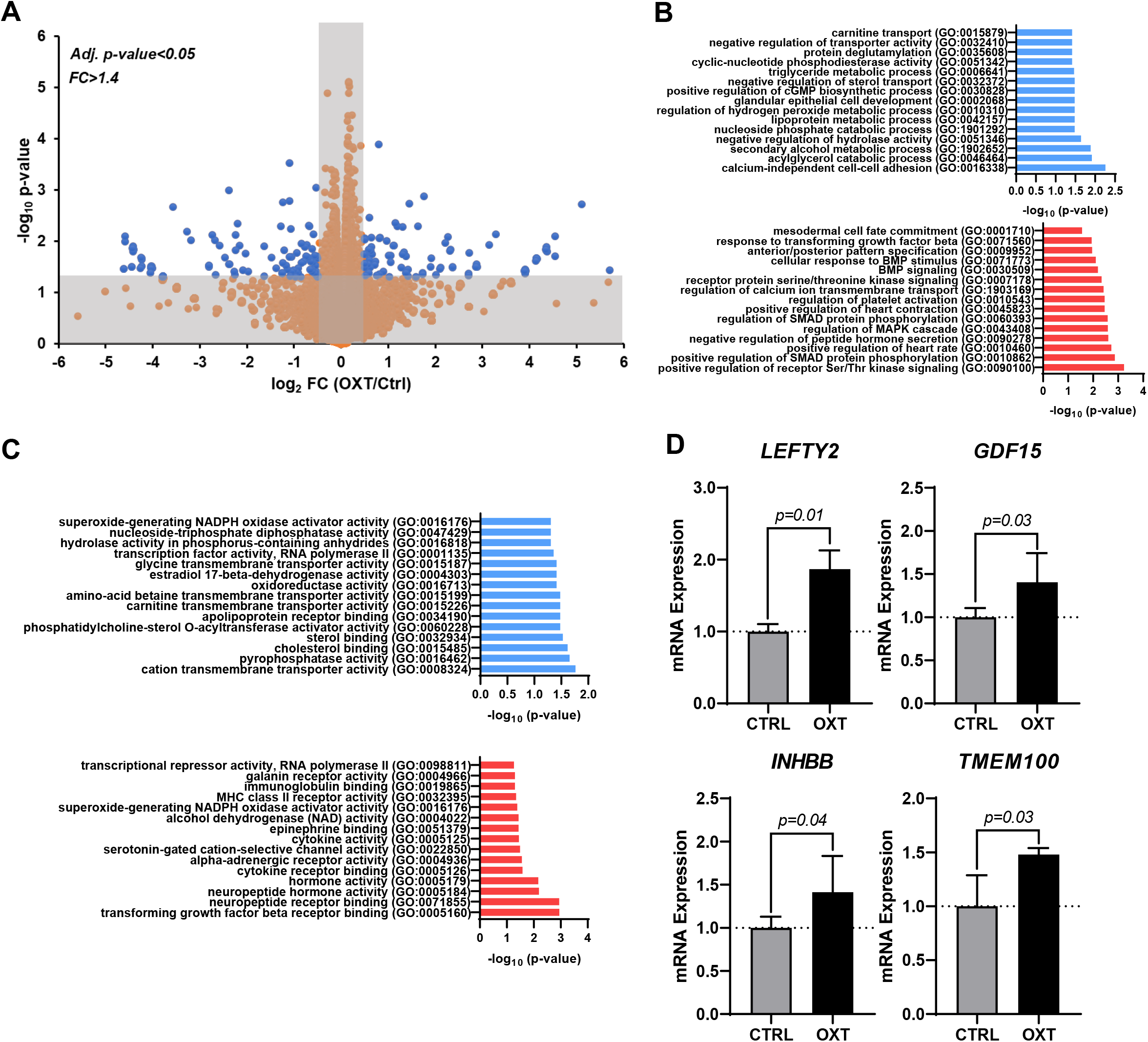
Transcriptome analysis of oxytocin-treated human epicardial cells. **A)** Volcano plot showing differentially expressed genes between control and OXT-treated hEpiCs, as determined by RNA sequencing; Blue dots correspond to genes with a fold change ≥1.4 and a p-value ≤0.05. **B-C)** Gene ontology analysis showing upregulated and downregulated biological processes (B) and molecular functions (C) after oxytocin exposure in hEpiCs. **D)** Relative mRNA expression of ligands for the TGF-β pathway and activators of BMP signaling in control and OXT-treated hEpiCs; n=3 per condition.

### Oxytocin signaling plays a role in the formation and migration of the proepicardial organ (PEO) during heart development

Our data suggest that OXT might play an important and overlooked role in epicardial function. We hypothesized that this may be reflected during heart development too. We therefore sought to determine if oxtr (the zebrafish ortholog to mammalian OXTR) inhibition would have deleterious effects on epicardium formation during heart development. The epicardium derives from an extra-cardiac cell cluster originating in the splanchnic mesoderm called the proepicardial organ (PEO). Once the PEO forms, it contributes new cells to the developing heart by migration into the epicardial and subepicardial spaces. These cells undergo EMT to form EpiPCs, which go on to differentiate into mature EpiCs and other cardiac cell types (Quijada et al., 2020; Simões & Riley, 2018). We bred double transgenic myl7-DsRed2, tcf21-nls-eGFP zebrafish embryos with a dsRed2 reporter in cardiomyocytes and an enhanced green fluorescent protein (EGFP) tag attached to a nuclear localization signal for *tcf21*, a well-established EpiPC marker (Kikuchi et al., 2010; J. Wang et al., 2011). This method allowed us to visualize the development of the epicardium and myocardium in real time via fluorescence microscopy. We dissolved different concentrations of L-368,899, a non-peptide OXTR antagonist, into zebrafish embryo medium and imaged epicardium formation at different time points (3, 5 and 7 dpf), quantifying the number of tcf-21 positive (green) cells covering the ventricular myocardium (red). L-368,899 induced a dose-dependent and statistically significant decrease in the number of epicardial cells covering the ventricle shortly after PEO formation, at 3 dpf (**Figure 5A, B**). As time passed, this difference intensified and manifested as a delay in epicardial formation at 5 and 7 dpf (**Figure 5C-F**). We also found that oxtr inhibition adversely affects formation of the atrial epicardium, as tcf21^+^ nuclei counts on the atrium were modestly (~15-30%) lower after drug treatment too (**Supplementary Figure 2A, B**). We confirmed the above findings by repeating this experiment with atosiban, a competitive OXTR antagonist (**Supplementary Figure 2C, D**). We concluded that oxytocin receptor inhibition significantly disrupts the formation of the epicardium in zebrafish. This delay could be due to defects in proepicardial cell migration to the surface of the developing heart, as it appears that the proepicardial cluster is farther from the myocardium in the atosiban-treated embryos than in control embryos (**Supplementary Figure 2C, D**). Alternatively, it could be due to a defect in epicardial cell proliferation, as both migration and proliferation of epicardial progenitors seem to be affected by OXT.

**Figure 5:**
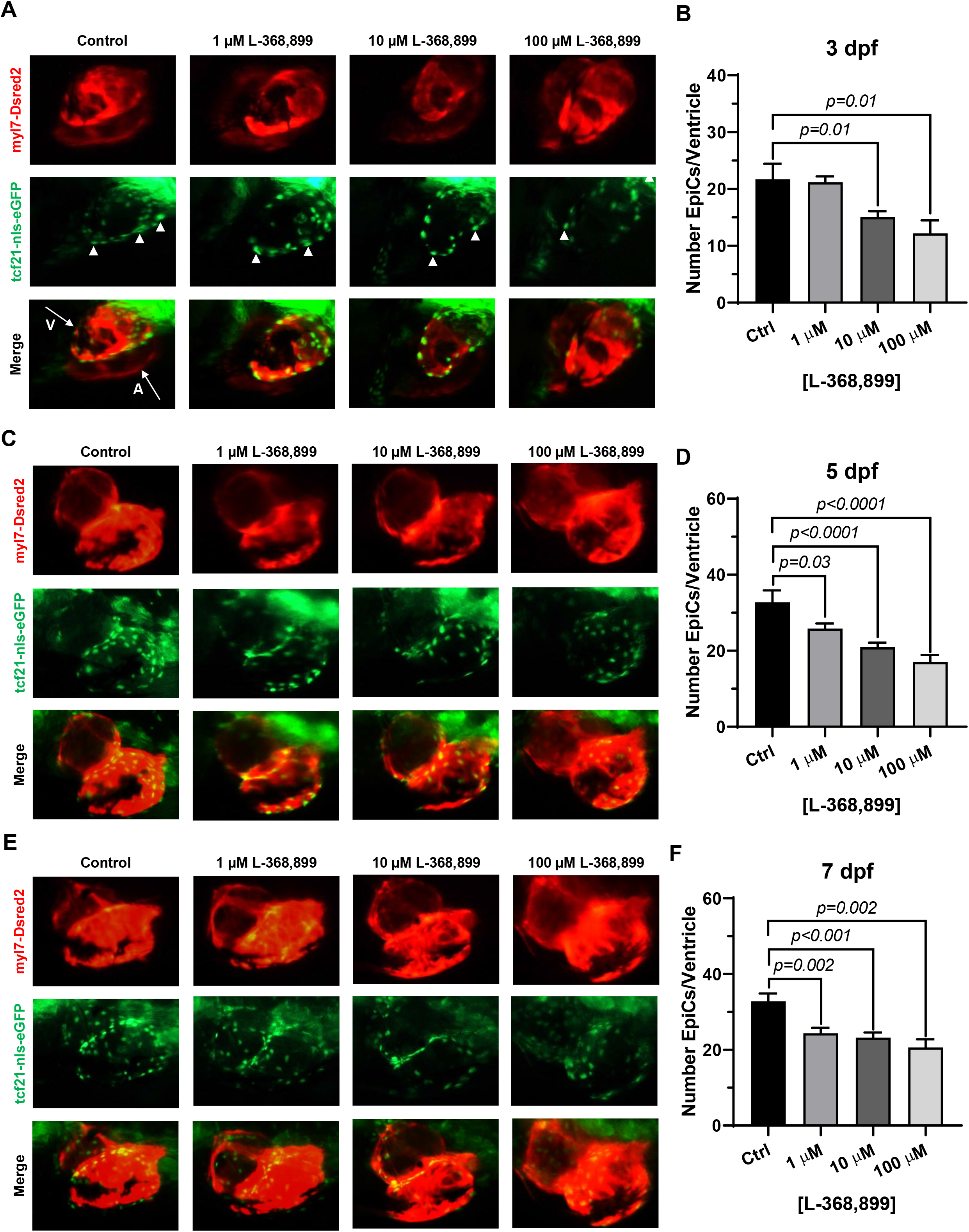
Oxytocin signaling is necessary for epicardium development in zebrafish. **A-F)** Fluorescent images (A, C, E) and epicardial cell counts per ventricle (B, D, F) of developing zebrafish embryos at 3, 5, and 7 dpf treated with different concentrations of L-368,899. In (A), (C), and (E) proepicardial and epicardial cell nuclei are labeled with GFP (tcf21-nls-eGFP, arrowheads), myocardium is labeled with DsRed2 (red); n≥5 embryos per condition for each time point; A: Atrium, dpf: Days Post-Fertilization, V: Ventricle.

### Oxytocin signaling is necessary for heart regeneration *in vivo*

The zebrafish is a powerful naturally regenerating organism popular for heart regeneration studies (Aguirre et al., 2014; Jopling et al., 2010; Poss et al., 2002). We decided to test if the role OXT plays in epicardial activation was conserved during zebrafish heart regeneration using a well-established cardiac cryoinjury model (González-Rosa et al., 2011; González-Rosa & Mercader, 2012) (**Figure 6A**). After cryoinjury, we observed an 18-fold increase in *oxt* mRNA levels in zebrafish brains at 3 dpi (days-post injury) and lasting until at least 7 dpi (**Figure 6B**), suggesting that a burst of oxytocin is released from the brain into the bloodstream after cardiac injury over a sustained period of time. To determine the functional significance of this oxytocin burst, we injected the specific OXTR inhibitor L-368,899 directly into the thoracic cavity of the fish after cryoinjury right after the procedure and every other day afterwards. The size of the injured myocardial area was assessed by Masson’s trichrome staining after injury in control and L-368,899 treated hearts. The differences in scar size between control and treated animals increased as time passed after cryoinjury. Inhibitor-treated animals displayed a modest increase in injured myocardium (~1.25-1.5 fold) at 3 and 7 dpi and a ~2-fold statistically significant increase at 14 dpi (**Figure 6C, D**). In addition, injured hearts treated with L-368,899 had more myocardial fibrosis accumulation than control hearts at 14 dpi (**Figure 6E**). Taken together, these data suggested that oxytocin signaling is necessary for proper heart regeneration. Two typical physiological responses observed in zebrafish heart regeneration are cardiomyocyte proliferation and endothelial revascularization adjacent to the injured area (González-Rosa et al., 2017; Quijada et al., 2020; Smits et al., 2018; Lavine et al., 2005; Li et al., 2011). Therefore, we collected control and treated hearts at 3 dpi and co-stained for PCNA (a commonly used marker of cell proliferation in zebrafish heart regeneration studies) (Aguirre et al., 2014; Jopling et al., 2010) and TNNT2, a specific cardiomyocyte marker. We observed a significant >2-fold decrease in cardiomyocyte proliferation at the infarct border zone after L-368,899 administration (**Figure 6F, G**). We also conducted cryoinjuries in fli1-eGFP zebrafish, a transgenic strain that specifically labels endothelial and endocardial cells with GFP (Lawson & Weinstein, 2002). We found that GFP fluorescence in the cryoinjured area was >1.5-fold higher in control fish compared to treated animals, suggesting that endothelialization and revascularization of the wound are slowed by oxtr inhibition (**Figure 6H, I**). The deleterious effects of L-368,899 on regeneration are very likely indirect, as cardiomyocytes and endothelial cells express very low or undetectable levels of oxtr (data not shown). In this context, activated epicardial cells have been shown before to promote revascularization and cardiomyocyte proliferation by transdifferentiation and secretion of paracrine factors (Lavine et al., 2005; Li et al., 2011).

**Figure 6:**
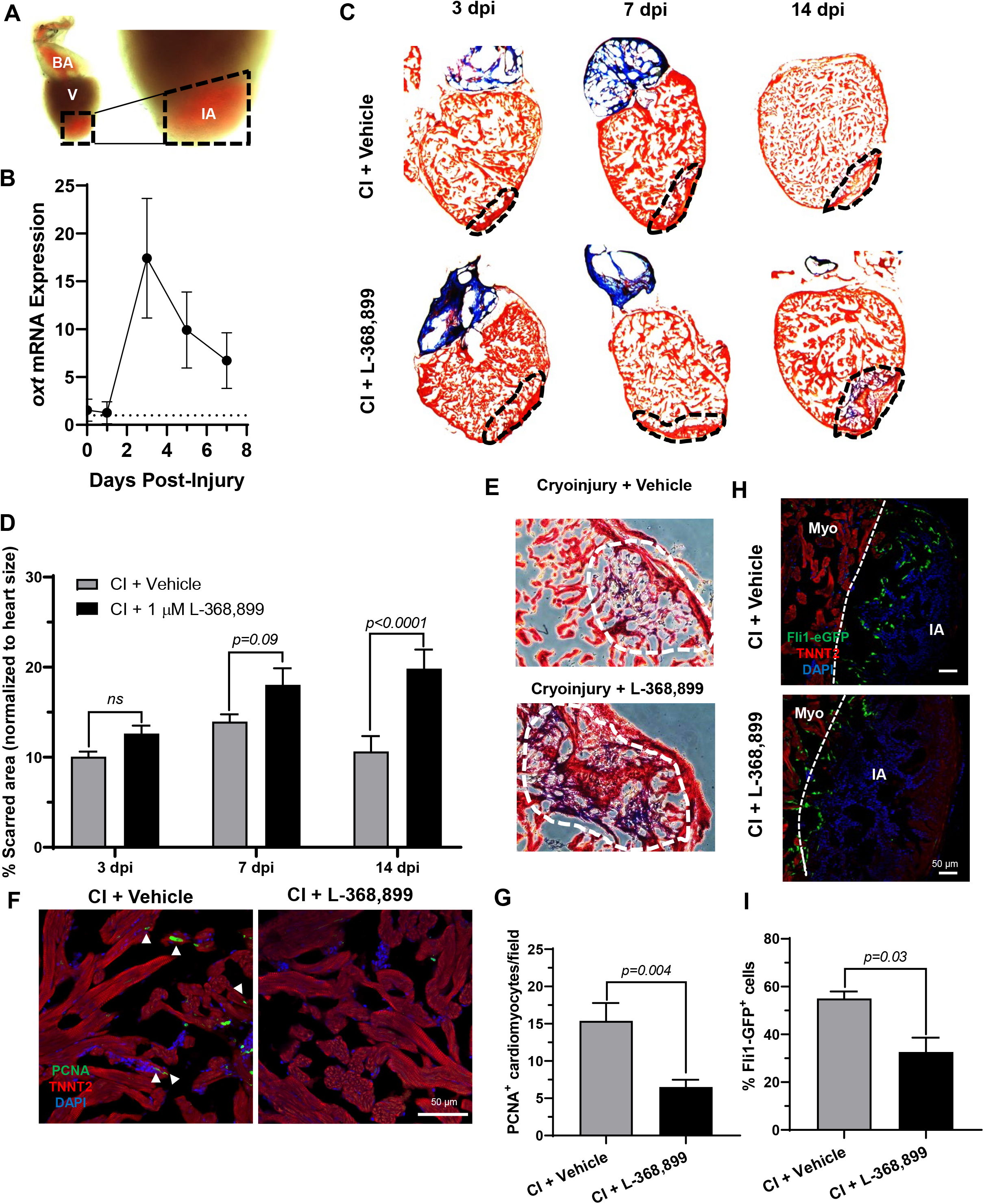
Oxytocin signaling is necessary for proper heart regeneration in zebrafish. **A)** Example of a freshly cryoinjured zebrafish heart, dashed lines demarcate injured area. **B)** Time course qRT-PCR data for *oxt* in zebrafish brains after cardiac cryoinjury; n=3 per time point. **C-D)** Masson’s trichrome staining (C) and quantification (D) of cryoinjured zebrafish hearts at 3, 7, and 14 dpi treated with and without 1 μM L-368,899. In (C), myocardium is stained red, collagen is stained blue, dashed lines demarcate injured area, which is quantified in (D); n=3-5 hearts per time point. **E)** Representative images of cryoinjured zebrafish myocardium (red) at 14 dpi showing more fibrosis accumulation (blue) after OXTR inhibition. Dashed lines demarcate fibrotic area. **F-G)** Confocal immunofluorescent images (F) and quantification (G) of proliferating cardiomyocytes in cryoinjured zebrafish hearts at 3 dpi adjacent to injured areas. In (F), cardiomyocytes are labeled with TNNT2 (red), proliferating cells (arrowheads) with PCNA (green), nuclei with DAPI (blue); Left images are low magnification (scale bar: 50 μm), right images are high magnification of myocardium immediately adjacent to the injured area (scale bar: 25 μm); n=8 images per condition. **H-I)** Confocal immunofluorescent images (H) and quantification (I) of *fli1-eGFP* transgenic zebrafish at 3 dpi showing endothelial revascularization in cryoinjured and L-368,899-treated hearts. In (H), endothelial and endocardial cells are labeled with GFP (green), cardiomyocytes with TNNT2 (red), nuclei with DAPI (blue); n=3-4 images per condition, scale bar: 50 μm; BA: Bulbus Arteriosus, CI: Cryoinjury, CM: Cardiomyocyte, dpi: Days Post-Injury, IA: Injured Area, Myo: Myocardium, V: Ventricle.

Next, we decided to explore how blocking oxt affected epicardial progenitor cell activation *in vivo*. Because one hallmark of epicardial activation after cardiac injury *in vivo* is upregulation of WT1 levels (Quijada et al., 2020; Smits et al., 2018), we next collected cryoinjured hearts to quantify its expression. We found that *wt1b* increased significantly after cryoinjury, with peak mRNA levels at 3 dpi (**Figure 7A**) and expression patterns very consistent with the brain release of *oxt* (**Figure 6B**). In addition, immunofluorescence using transgenic tcf21-nls-eGFP fish showed an expansion and migration of epicardial progenitors into the subepicardial and myocardial layers of the heart, but these effects were abolished in L-368,899 treated animals (**Figure 7B-D**). We also found that L-368,899 treated zebrafish hearts had significantly lower expression of *wt1b* and *tcf21* compared to control hearts (**Figure 7E**) and decreased expression of EMT markers, as revealed by *snai1a* and *snai2* mRNA levels (~40-50% reduction) (**Figure 7F**). Thus, expansion of the subepicardial area and migration of WT1+ EpiPCs into the myocardium, two hallmark of zebrafish heart regeneration (González-Rosa et al., 2017; Quijada et al., 2020; Smits et al., 2018), were gravely impaired upon oxytocin signaling inhibition. We concluded that oxtr inhibition prevents robust activation of the epicardium and its progenitor cell populations after injury, leading to an impaired regenerative response.

**Figure 7:**
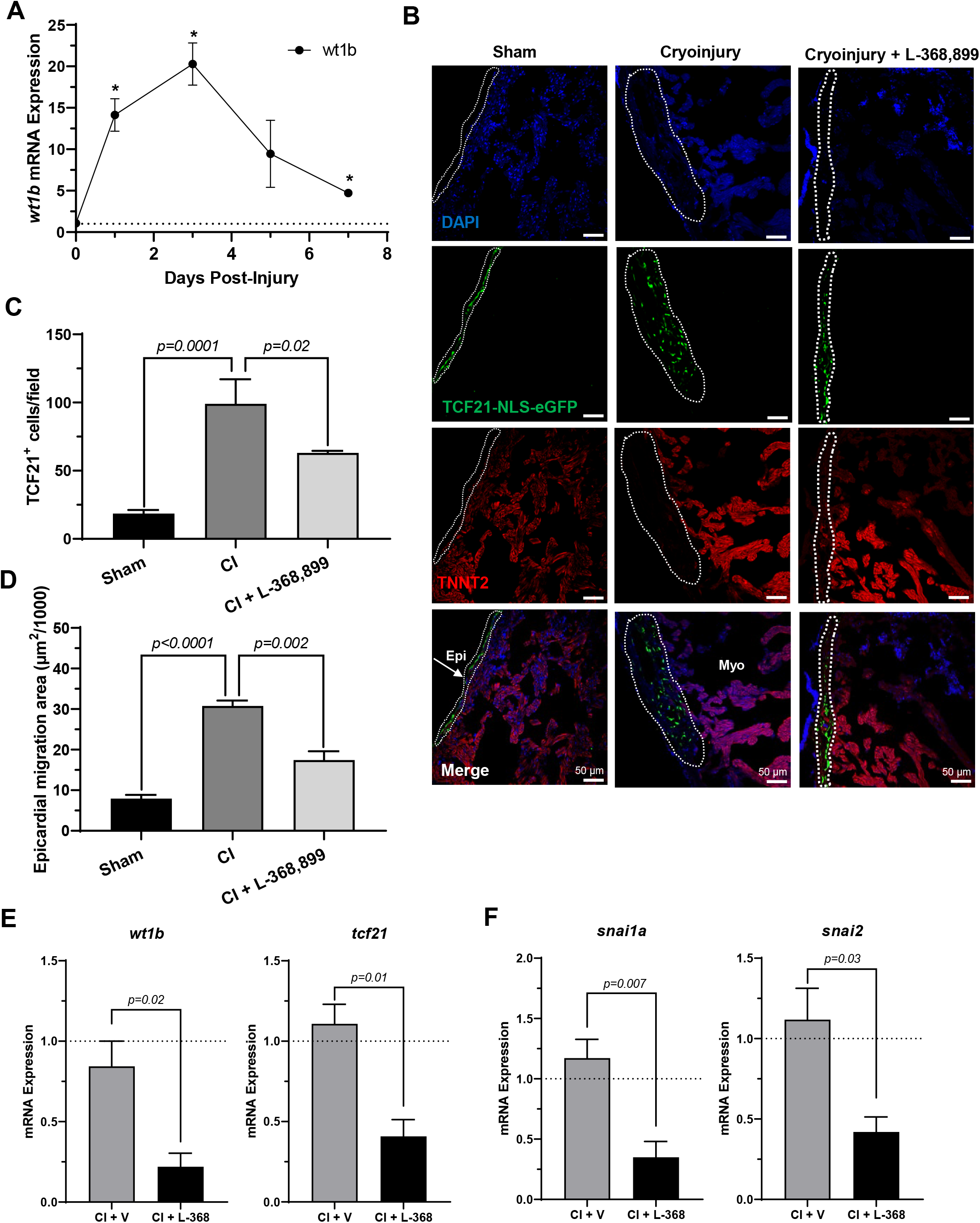
Inhibition of oxytocin signaling prevents epicardial activation after cardiac injury. **A)** Time course qRT-PCR data for *wt1b* in zebrafish hearts after cardiac cryoinjury; *P<0.05 versus sham operated heart; n=3 per time point. **B-D)** Confocal immunofluorescent images (B) and quantification (C-D) of *tcf21-nls-eGFP* transgenic zebrafish at 3 dpi showing epicardial activation in sham, cryoinjured, and L-368,899-treated hearts. In (B), epicardial cell nuclei are labeled with GFP (green), cardiomyocytes are labeled with TNNT2 (red), nuclei are labeled with DAPI (blue); n=3-5 images per condition, scale bar: 50 μm. **E-F)** qRT-PCR data for *wt1b* and *tcf21* (E) and *snai1a* and *snai2* (F) in cryoinjured zebrafish hearts at 3 dpi treated with and without 1 μM L-368,899 (L-368); n=4-5 per condition; CI: Cryoinjury, dpi: Days Post-Injury, Epi: Epicardium, Myo: Myocardium, SE: Subepicardium, V: Vehicle.

## DISCUSSION

Oxytocin is a hypothalamic neuroendocrine hormone best known for its functions in parturition, lactation, and social bonding. However, it also plays other less known physiological roles, including in the cardiovascular system, where it lowers blood pressure, induces negative inotropic and chronotropic effects, and serves as an anti-inflammatory and anti-oxidant (Garrott et al., 2017; Gutkowska et al., 2014; Jankowski et al., 2020). Many of these actions are carried out by OXT-mediated release of atrial natriuretic peptide (ANP) and nitric oxide (NO) (Jankowski et al., 2020), which have well-characterized cardioprotective effects (Jones & Bolli, 2006; Nishikimi et al., 2006) and may themselves contribute to heart regeneration (Kook et al., 2003; Rochon et al., 2020). Notably, OXT also induces differentiation of cardiomyocytes from embryonic stem cells (Paquin et al., 2002). Recent studies suggest that damage to neuroendocrine structures, such as the hypothalamus, strongly inhibits endogenous regenerative processes in vertebrates (Zhang et al., 2018), suggesting that critical pro-regenerative factors are released from this brain region. In addition, acute myocardial infarction activates OXT-releasing neurons in the paraventricular nucleus of the rat hypothalamus (Roy et al., 2019). The above studies, combined with ample evidence to suggest that epicardial activation can be primed by specific signaling factors, such as thymosin β4 (Smart et al., 2007, 2011), suggest that oxytocin may be the critical factor in achieving sufficient EpiPC activation and differentiation to regenerate the lost myocardium after cardiac injury.

Here, we have shown that OXT induces a pro-regenerative phenotype *in vitro* in human epicardial cells. When mature-like hEpiCs were exposed to OXT, they increased their proliferation rates (**Figure 2B-D**) and adopted a stem cell-like gene expression profile (**Figure 2F**), two hallmarks of epicardial activation. Notably, oxytocin induced an even greater proliferative response than thymosin β4 (**Figure 2E**). We found that OXT acted through its receptor to exert its effects as the increases in hEpiC proliferation and activation that were seen in scrambled cells were completely abolished in *OXTR* knockdown epicardial cells (**Figure 3D-G**). Although OXT is also able to bind to AVP receptors with lower affinity (Gimpl & Fahrenholz, 2001), it does not appear as if this signaling pathway plays a role in our model, as hEpiCs do not express these receptors. In order to further demonstrate the importance of oxytocin signaling to heart regeneration, we extended our experiment to zebrafish, one of the most powerful naturally regenerating animal models known (Beffagna, 2019; Jopling et al., 2010; Poss et al., 2002). Several seminal studies of the last ~20 years have shown that the epicardium plays a crucial role in zebrafish heart regeneration (Cao et al., 2016; Cao & Poss, 2018; Kikuchi et al., 2011; Lepilina et al., 2006), therefore it is the perfect model system with which to expand our studies *in vivo*. We demonstrated that pharmacological inhibition of OXT signaling significantly slows cardiac regeneration after cryoinjury, as seen by increased fibrosis accumulation and myocardial injury area when zebrafish are treated with L-368,899 (**Figure 6C-E**). In addition, this compound decreases activation and proliferation of CMs as well as revascularization of the wound (**Figure 6F-I**). OXTR inhibition also prevents the proliferative response of epicardial progenitor cells that is normally seen after cryoinjury (**Figure 7B, C**). It is certainly possible that the anti-proliferative effects L-368,899 treatment has on the epicardium ultimately cause these effects in the other cell types, possibly through indirect pathways related to epicardial secretion of pro-regenerative molecules (Lavine et al., 2005; Li et al., 2011). Future studies will further explore the interaction of EpiPCs with CMs, endothelial cells, and other cardiac cells, such as cardiac fibroblasts and immune cells. Our data also suggest that OXT signaling is required for proper epicardial activation, as L-368,899 treatment decreases the cardiac expression of several genes that are involved in EpiPC expansion and EMT (**Figure 7E, F**). Our hypothesis is that oxytocin is released from the brain into the bloodstream after cardiac injury to facilitate epicardial activation and heart regeneration. We showed that *oxt* mRNA levels increase ~20-fold in zebrafish brains 3 days after cardiac cryoinjury (**Figure 6B**), with a corresponding increase seen in *wt1b* expression in the injured hearts (**Figure 7A**).

In conclusion, we show here that oxytocin is an important activator of EpiPCs and is necessary for heart regeneration and proper epicardium development. We demonstrate that OXT induces reprogramming of human epicardial cells into an epicardial progenitor state by activating the OXTR. These effects are mediated through the TGF-β pathway and its associated downstream effectors. Overall, our results establish a previously uncharacterized link between OXT release after cardiac injury and heart regeneration and provide evidence to support the notion that cardiac regeneration is, at least partially, under neuroendocrine control. These findings offer exciting translational potential, as oxytocin or one of its agonists could be used in the clinic to aid in the recovery from severe cardiac events, such as an MI, and prevent progression to heart failure in the future.

## AUTHOR CONTRIBUTIONS

A.H.W. and A.A. designed all experiments and conceptualized the work. A.H.W. performed all experiments, acquired images, and analyzed data. A.R.H. performed cell culture, zebrafish husbandry, and experiments. Y.R.L.-I. acquired *in vitro* data and confocal images. M.D.D. and A.L.M. acquired and analyzed molecular biology data. M.V. performed zebrafish experiments. A.H.W. prepared all figures, and A.H.W. and A.A. wrote the manuscript. A.A. supervised all the work.

## ACKNOWLEDGEMENTS

We wish to thank the MSU Genomics Core for RNA sequencing support and Dr. William Jackson at the MSU Department of Pharmacology/Toxicology for access to confocal microscopes. We also want to thank all members of the Aguirre lab for helpful comments and discussions. Work in Dr. Aguirre’s laboratory is supported by the National Heart, Lung, and Blood Institute of the National Institutes of Health under award numbers K01HL135464 and R01HL151505, by the American Heart Association under award number 19IPLOI34660342, and by the Spectrum-MSU Foundation.

