## Supplementary data for "Oxytocin promotes epicardial cell activation and heart regeneration after cardiac injury"

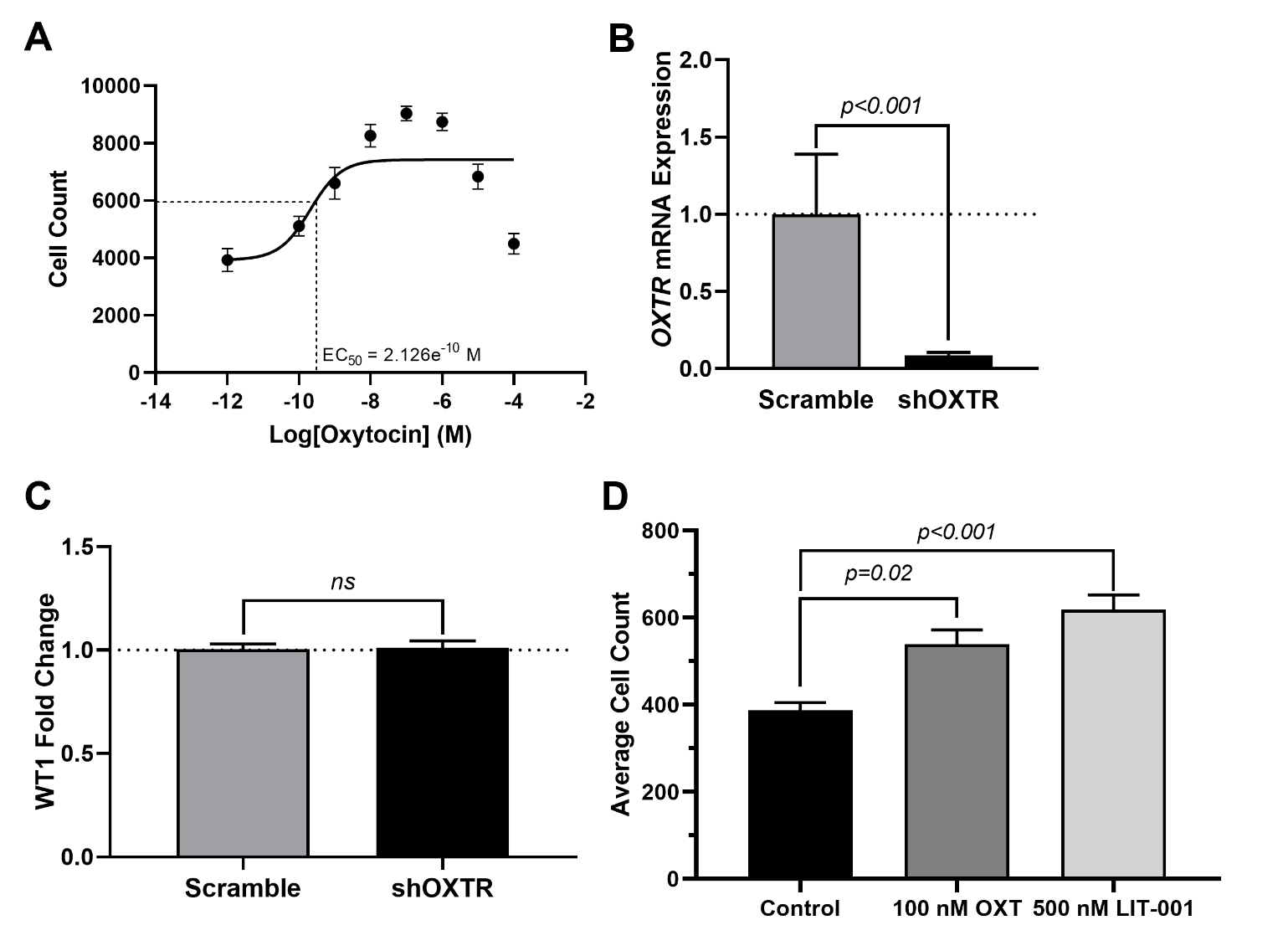

**Supplementary Figure 1**. **A)** Oxytocin dose-response curve for hEpiCs over the course of 5 days. **B)** qRT-PCR for scrambled and OXTR knockdown for hEpiCs demonstrating a ~85% knockdown of OXTR. **C)** Efficiency of hiPSC differentiation into hEpiCs in scrambled and shOXTR cell lines by qRT-PCR for WT1 expression at day 25 of differentiation. **D)** Proliferation effects of OXT and the specific OXTR agonist LIT-001 on hEpiCs. N=3-5 independent experiments in all cases.

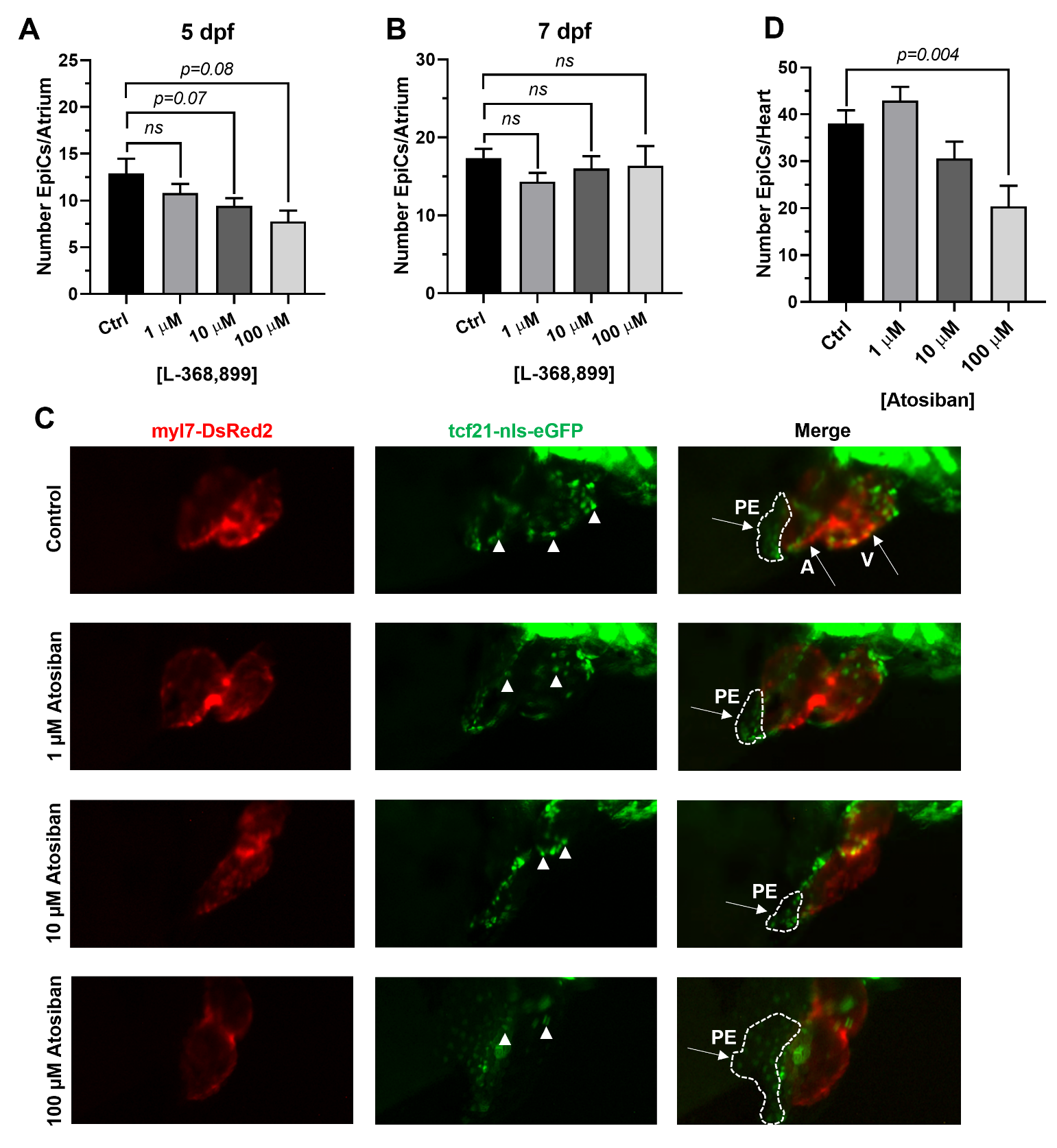

**Supplementary Figure 2**. Epicardial cell counts per atrium in developing double transgenic myl7-DsRed2; tcf21-nls-eGFP zebrafish embryos at 5 (**A**) and 7 dpf (**B**) treated with different concentrations of the small molecule oxytocin receptor inhibitor L-368,899. **C,** Fluorescent micrographs of developing myl7-DsRed2; tcf21-nls-eGFP embryos at 3 dpf treated with different concentrations of atosiban, a peptide oxytocin receptor inhibitor. D) Quantification of the atosiban effects on epicardial cell migration at 3 dpf. Epicardial tissues appear green (arrowheads), while cardiomyocytes appear red. Dashed lines demarcate the proepicardial organ; n≥8 embryos per condition; A: Atrium, PE: Proepicardium, V: Ventricle.
