## Supplementary Table 1 for "Oxytocin promotes epicardial cell activation and heart regeneration after cardiac injury"

**Supplementary Table 1**. Neuroendocrine Candidate List for Screening in hEpiCs

| **Name** | **Abbreviation** | **Site of Production** | **Site of Release** | **Regenerative Effect** | **References** |
| --- | --- | --- | --- | --- | --- |
| Adrenocorticotropic Hormone | ACTH | Anterior Pituitary | Anterior Pituitary | Cortisol inhibits heart and tailfin regeneration in zebrafish | Sallin and Jazwinska, 2016; Hartig et al., 2016 |
| Brain-Derived Neurotrophic Factor | BDNF | Many Brain Regions | Many Brain Regions | Stimulates proliferation of neural stem cells (by activating Wnt signaling) | Chen et al., 2013 |
| Growth Hormone | GH | Anterior Pituitary | Anterior Pituitary | Full limb regeneration in hypophysectimized newts | Landesman and Hessler, 1981; Landesman and Copeland, 1988 |
| Growth Hormone-Releasing Hormone | GHRH | Hypothalamus | Hypothalamus | GHRH agonist increases mitosis and decreases infarct size in rats after MI | Kanashiro-Takeuchi et al., 2012 |
| Melanin-Concentrating Hormone | MCH | Hypothalamus | Hypothalamus | Stimulates GH secretion and may promote pancreatic islet regeneration | Segal-Lieberman et al., 2006; Pissios et al., 2007 |
| Neuropeptide Y | NPY | Hypothalamus | Hypothalamus | Promotes mitosis in vascular smooth muscle cell cultures | Pons et al., 2003 |
| Oxytocin | OXT | Hypothalamus | Posterior Pituitary | Induces CM differentiation and stimulates muscle regeneration in mice | Paquin et al., 2002; Jankowski et al., 2004; Elabd et al., 2014 |
| Pituitary Adenylate Cyclase Activating Polypeptide | PACAP | Pituitary Gland | Pituitary Gland | Stimulates axonal regeneration after spinal cord injury in mice | Tsuchida et al., 2014 |
| Pro-Opiomelanocortin | POMC | Hypo./Pituitary | Cleaved to Peptides | Cleavage products have numerous regenerative effects | N/A |
| Prolactin | PRL | Anterior Pituitary | Anterior Pituitary | Promotes tail regeneration in newts and regulates liver regeneration in mice | Liversage et al., 1984; Moreno-Carranza et al., 2013 |
| Somatostatin | GHIH | Hypothalamus | Hypothalamus | Inhibits limb and tail regeneration in newts | Vethamany-Globus et al., 1977 |
| Thyroid Stimulating Hormone | TSH | Anterior Pituitary | Anterior Pituitary | Thyroid hormone (TH) surge in mice at PND 15 causes CM proliferation | Naqvi et al., 2014 |
| Thyrotropin-Releasing Hormone | TRH | Hypothalamus | Hypothalamus | Stimulates epidermal regeneration in tadpole and human skin | Meier et al., 2013 |
| α-Melanocyte-Stimulating Hormone | α-MSH | POMC/ACTH Cleavage | Hypo./Brainstem | Binds to Mc4r, which is required for tadpole limb regeneration | Zhang et al., 2018 |
| β-Endorphin | β-End | POMC/ACTH Cleavage | Hypo./Pituitary | Stimulates forelimb regeneration in hypophysectimized newts | Morley and Ensor, 1986 |
